# Base Composition, Speciation, and Why the Mitochondrial Barcode Precisely Classifies

**DOI:** 10.1101/116814

**Authors:** Donald R. Forsdyke

## Abstract

While its mechanism and biological significance are unknown, the utility of a short mitochondrial DNA sequence as a “barcode” providing accurate species identification has revolutionized the classification of organisms. Since highest accuracy was achieved with recently diverged species, hopes were raised that barcodes would throw light on the speciation process. Indeed, a failure of a maternally-donated, rapidly mutating, mitochondrial genome to coadapt its gene products with those of a paternally-donated nuclear genome could result in developmental failure, thus creating a post-zygotic barrier leading to reproductive isolation and sympatric branching into independent species. However, the barcode itself encodes a highly conserved, species-invariant, protein, and the discriminatory power resides in the non-amino acid specific bases of synonymous codons. It is here shown how the latter could register changes in the oligonucleotide frequencies of nuclear DNA that, when they fail to match in pairing meiotic chromosomes, could reproductively isolate the parents (whose hybrid is sterile) so launching a primary divergence into two species. It is proposed that, while not itself contributing to speciation, the barcode sequence provides an index of the nuclear DNA oligonucleotide frequencies that drive speciation.

## Introduction

The discovery that a short mitochondrial DNA (mtDNA) sequence could serve as a “barcode” providing rapid and accurate species identification (Hebert et al. 2003a) has revolutionized the classification of organisms. I here consider the mechanism and significance for speciation of the barcode’s classificatory (taxonomic) effectiveness. The paper has four parts. The first considers whether the barcode operates at the initiation of species divergence and, if not, when after that divergence the barcode emerges as a species attribute. The second discusses another, apparently independent, species attribute, the base composition of DNA. This internal “genome phenotype” is a partial driver of the amino acid composition of the proteins that, directly or indirectly, have generated the characters (conventional phenotype) upon which classification has traditionally depended. The third part shows how genetic code degeneracy affords a distinction between genome and conventional phenotypes. Finally, the fourth part turns to mechanisms for the initiation of divergence of one species into two, and considers how information from oligonucleotide frequencies that relate to the genome phenotype could be transferred to mitochondrial DNA, where the barcode sequence would best register that phenotype and hence the species. It is concluded that such mitochondrial calibration is an incidental by-product of more fundamental evolutionary processes and is not itself involved in the initiation of species.

## 1. Species-Specific Barcode

### Classification by a Single Attribute

In keeping with accepted norms of evidence, those who classify organisms into species have valued multiple attributes. In popular parlance, “when I see a bird that walks like a duck and swims like a duck and quacks like a duck, I call that bird a duck.” That DNA sequences might likewise facilitate classification came as no surprise. Similarity between two organisms in few parts of long sequences indicates evolutionary distance; they are members of different species. Conversely, similarity between two organisms in many parts of long sequences indicates evolutionary closeness; they may even be members of the same species. This multiple attribute requirement (Altschul et al. 1990) was starkly challenged by the discovery of Canadian biologist Paul Hebert that a fragment of the genome of a minute cytoplasmic organelle could stand in for the entire nuclear genome. Ignoring the rest of an organism’s DNA, a mere 648 base mitochondrial DNA sequence (part of the COI gene) could act as a “species barcode” giving high precision species identification (Hebert et al. 2003a).

While the term “barcode” raised eyebrows, and problems still remain (especially with plants;Chase and Fay 2009), its accuracy is now clearly recognized. Yet its mechanism and biological significance remain a mystery. There are many 648 base sequences in genomes. Indeed, there are 4^648^ possible combinations of 648 bases. In principle any such sequence, or an even smaller one, could serve as a barcode. The mystery is why a segment, in a particular gene, in a mitochondrion, can be singled out for this role?

The involvement of mitochondria is less problematic. The higher mutation rate of mtDNA relative to nDNA (nuclear DNA) creates greater between-species DNA diversity in mitochondria. Hebert envisaged that *within* members of a species there is “a scouring mechanism that cleanses mitochondrial genomes of their diversity on a regular basis” (Strauss 2006). This permits discrimination between members of closely related species that have recently diverged from a common ancestor, and sometimes even reveals cryptic species; that is, members of what were formerly regarded, on morphological grounds, as a single species are actually members of two or more sibling species.

### Role in Speciation

Focused on the utility of his barcode as an aid to classification, Hebert held that the differences between species that made barcoding so effective had arisen *subsequent* to the initiation of the branching into independent, reproductively isolated, species. Whatever the mechanism of barcoding, it was dependent on a *prior* initiation of branching, and had played no role in that initiation. Yet its sensitivity in detecting recently or cryptically diverged species suggested a more active role in the speciation process. To this Herbert, hailed in some quarters as “the new Linnaeus,” replied that “it would be a bit presumptuous to say that little gene bit had played a key role in speciation. You could say it if you wanted to but not me. I have enough nails through my hands from critics to not want to go in that direction just now” (Strauss 2006).

Nevertheless the view grew that “the mitochondrial bioenergetics system should be recognized as a candidate key driver of speciation events, at least in animals” (Gershoni et al. 2009). More specifically, it was proposed that, “the DNA bar code does not merely track species — it could very well create them.” The assumption that barcode sequences “became unique after two species split from their common ancestor” was probably incorrect (Lane 2009). Indeed, without provision of a detailed mechanism, a role in speciation was boldly held to *explain* the barcode phenomenon (Hill 2016): “An explanation for why the sequence of a specific gene coincides so consistently with species boundaries is that the barcode gene, or the other mitochondrial genes to which it is linked, plays a role in the process of speciation.”

### The Barcode Gap and Divergence Time

In general, “sequence divergences are much larger among species than within species, and thus mtDNA genealogies generally capture the biological discontinuities recognized by taxonomists as species” (Hebert et al. 2004). Thus, barcode sequence differences between individuals of the same species are very low (much less than 1%), whereas the differences, even between individuals from closely related (allied or congeneric) species, are uniformly greater (2% and higher). This “barcode gap” facilitates precise classification.

However, there were exceptions. It was thought that rare interspecies differences below 2% were related to a brief evolutionary period that had elapsed since the first sparking of the divergence into two species. Thus, while 98% of congeneric species pairs showed more than 2% divergence, Hebert et al. (2003b) noted that: “About 1.9% of the congeneric pairs in groups with normal rates of mitochondrial evolution showed less than 2% divergence, probably often reflecting short histories of reproductive isolation.” In other words, the exceptions were, on an evolutionary time-scale, very young species: “COI diagnoses should not fail … unless species are unusually young” (Hebert et al. 2003a). While calling for “a newly detailed view of the origins of biological diversity,” these thoughts sustained the impression that Hebert regarded species initiation as occurring *prior* to whatever barcodes were registering.

Herbert acknowledged that the high mutation rate of mtDNA would determine that, long after the initial speciation event, there would eventually be a time when the mtDNA divergence rate would slow, since every viable mutation between two mitochondrial sequences would have occurred; back-mutations would reverse forward-mutations and then the forward-mutations could recur, but no further net, time-dependent, divergence was likely. Thus: “Obviously, the analysis of rapidly evolving gene regions or taxa will aid the diagnosis of lineages with brief histories of reproductive isolation, while the reverse will be true for rate decelerated genes or species” (Hebert et al. 2003a). However, albeit limited, he later noted that barcodes might still play some role in evaluating relationships between well-established species: “Although mutational saturation limits the value of COI barcode sequences for the independent resolution of deep-level phylogenetic relationships, barcode data are currently being incorporated into several ‘tree of life’ projects” (Ward et al. 2009).

As for the barcode mechanism, at the outset it was speculated that the within-species scouring or pruning that stabilizes the barcode - so generating high species specificity - somehow involved *coadaptation* of a cell’s mtDNA and nDNA (Hebert et al. 2003b):

> The clear delineation of most congeneric species pairs indicates a surprising ferocity of lineage pruning. The restricted levels of intraspecific mitochondrial divergence that result from this pruning are critical to taxon diagnosis and they may have a simple explanation. … there is growing evidence that mitochondrial and nuclear genomes are linked in a *pas de deux* of surprising intimacy.

Beyond this Hebert would not go.

## 2. Base Composition

### Base Composition as a Species Attribute

Irrespective of actual sequence, it has long been known that the frequency of the mononucleotide bases G and C (written as GC%) is a species correlate. The “GC rule” of Chargaff (1951)is that the ratio of (G + C) to the total bases (A + T + G + C) tends to be constant in a particular species, but varies between species. Sueoka (1961)further pointed out that for individual “strains” of *Tetrahymena* (ciliated unicellular organisms), GC% tends to be uniform *throughout* the genome:

> If one compares the distribution of DNA molecules of *Tetrahymena* strains of different mean GC contents, it is clear that the difference in mean values is due to a rather uniform difference of GC content in individual molecules. In other words, assuming that strains of *Tetrahymena* have a common phylogenetic origin, when the GC content of DNA of a particular strain changes, all the molecules undergo increases or decreases of GC pairs in similar amounts. This result is consistent with the idea that the base composition is rather uniform not only among DNA molecules of an organism, but also with respect to different parts of a given molecule.

Of high importance for present considerations, Sueoka also found a link between GC% and reproductive isolation:

> DNA base composition is a reflection of phylogenetic relationship. Furthermore, it is evident that those strains which mate with one another (i.e. strains within the same ‘variety’) have similar base compositions. Thus strains of variety 1 …, which are freely intercrossed, have similar mean GC content.

Whether Sueoka’s “strains” or “varieties” were in fact “species” is uncertain. But uniformity of GC% is now known to apply to a wide variety of species. Each species can be considered to have a specific value, but there is often a distribution about this value that reflects genome fine-sectoring into regions of slightly lower or higher GC%. Since GC% is a pervasive genome-wide characteristic, there is sometimes no need to sequence an entire genome to measure its value. Indeed, short sequence segments may suffice (Zavala et al. 2005; Fournier et al. 2006). However, GC% does not have the discriminatory power of the bar code.

### Base Composition as Internal Driver of Amino Acid Composition

Sueoka (1961) also observed that the amino acid composition of proteins can be influenced, not only by the demands of the environment (natural selection) on the proteins, but also by the GC% of the genome encoding those proteins. From the genetic code it can be correctly inferred that AT-rich genomes tend to have more codons for certain amino acids (Phe, Tyr, Met, Ile, Asn, Lys), and GC-rich genomes tend to have more codons for another set of amino acids (Gly, Ala, Arg, Pro). The relative abundance of these two sets in an organism’s proteins correlates well with its GC%. Conversely, with a knowledge of an organism’s base composition, one can predict its relative amino acid composition (Singer and Hickey 2000). Thus, shifts in genome (nucleotide) and proteome (amino acid) compositions are coupled. However, DNA base compositions generally show much greater variation between species than amino acid compositions. Crick (1959)was puzzled:

> This large variation of DNA composition is very unexpected. The abundance of the various amino acids does not, as far as we know, vary much from organism to organism; leucine is always common, methionine usually rather rare. … the large variation reported for DNA needs some special explanation.

That “special explanation” seems now at hand. Brbić et al (2015) report that species oligonucleotide frequencies can predict species amino acid compositions with even higher accuracy than species mononucleotide frequencies (base compositions); only 9% of the amino acid variation between species remains unexplained. They conclude that “evolutionary shifts in overall amino acid composition appear to occur almost exclusively through factors shaping the global oligonucleotide content of the genome.” It seems that oligonucleotide frequency has more room to manoeuvre than amino acid frequency. It is *oligonucleotide composition* (frequency of certain oligonucleotides), which, quite distinctly from conventional, *externally-imposed,* natural selection, responds to unnamed “factors,” so acting as an *internal* driving force that independently affects amino acid compositions.

### Oligonucleotide Composition and Species Identification

A need for a formal distinction between external and internal factors has been long recognized. The terms “active” and “passive” (Schaap 1971) were followed by “organismal phenotype” and “intragenomic phenotype” (Orgel et al. 1980), which were later superceded by mere “phenotype” – the traditional term meaning something upon which classical Darwinian natural selection might sometimes act – and “genome phenotype” (Bernardi and Bernardi 1986).

That the genomic shaping “factors” of Brbić et al. (2015) might relate to species became clearer when oligonucleotide frequency was found to support “metagenomic” species identification for assembling phylogenetic trees or genome sequences. In the latter case there is reversal of usual sequencing methodology – prepare DNA fragments from an identified organism, sequence them, then align overlapping fragments to obtain its full length genome sequence. Instead, DNA is first prepared from a source that contains numerous organisms. Then the fragments are sequenced. Individual species within the sample are *identified on the basis of their distinctive oligonucleotide frequencies.* In this manner one DNA sample can yield the sequences of numerous entire genomes, which only later are correlated with particular organisms (Dubinkina et al. 2016).

Thus, 648 base mtDNA barcodes and nDNA oligonucleotide frequencies *share* the property of being sensitively diagnostic of species. That this sensitivity might be greater than that provided by GC% alone had been deduced by Rogerson (1991), who used statistical approaches to taxonomic placement (species cluster and principle component analyses) that were similar to those that Richard Grantham had employed a decade earlier (see later):

> At first it would seem that the GC content of the DNA of the species might offer a reasonable explanation for their placement, as most species with low GC content tend to cluster together. However, species with higher GC content do not so cluster, and ordering the species by GC content … does not yield enough information to reproduce either the cluster diagram or the major components diagram.

## 3. Codons

### Codon Positions Reveal Phenotype Disconnect

A high GC% DNA segment is likely to contain oligonucleotides enriched in G and C – e.g. GCG, CGG, etc., rather than ATA, TAA, etc.. As discussed above, these oligonucleotides are likely to have had more input into GC% values than the converse (Rogerson 1991; Forsdyke 1995; Forsdyke and Bell 2004; Brbić et al. 2015). However, at this point I refer to a sequence’s GC% rather than to the oligonucleotide frequencies that relate to that GC%.

The mtDNA of some species has a low GC% (and hence a high level of A and T). In other species there is a high GC% (and hence a low level of A and T). As with nDNA, short sequences suffice for measurement of the GC% of mtDNA, which usually differs greatly from that of the corresponding nDNA. Min and Hickey (2007)showed for a wide range of organisms that there is a very close correlation between the GC% of COI barcode sequences and that of the corresponding mtDNAs. In other words, a barcode’s GC% and the barcode itself are species correlates. Thus, it might be suspected that, despite the difference from the nDNA’s GC%, a barcode’s GC% would somehow contribute to its species-discriminating power.

Indeed, Herbert and Gregory (2005) noted that whatever its GC% value, barcode accuracy is high. Thus: “Shifts in nucleotide composition of the mitochondrial genome … fail to impact the resolution of DNA barcoding, as evidenced by success in groups, such as birds, with high G+C composition and others, such as spiders, with extreme A+T bias.” Since *large* shifts in GC% values do *not* diminish a barcode’s resolving power, they are likely to be contributing positively to that power.

As we shall see, a barcode’s GC% value has two, largely independent, input sources. One relates to the encoded COI protein and is constrained by its needs. The other relates to the overall mtDNA GC% that seems constrained by the needs of the species. The difference is manifest in the structure of the barcode’s 216 codons that determine the amino acid sequence it encodes.

Either directly or indirectly, the anatomical and physiological characters of an organism (its phenotype) are determined by its proteins. Its conventional classification (phenotype-based) is a function of how much its proteins differ from those of other organisms. If a protein is highly variable across species, then it may assist species classification. If a protein is highly conserved across species then it is unlikely to assist species classification. The protein encoded by the COI gene is highly conserved (Pesole et al. 1999), so the aspect of barcode GC% that relates to this protein *cannot* contribute to the barcode’s discriminatory power.

The COI gene contains sequential sets of three bases (codons) that determine the nature and order of the amino acids in a cytochrome c oxidase subunit. The 648 base COI barcode segment encodes 216 amino acids. A high GC% bird mitochondrion synthesizes essentially the same subunit as a low GC% spider mitochondrion. This is because, as further discussed below, the shifts in the segment’s base composition between species mostly affect bases at codon positions that do *not* change the nature of the encoded amino acid. Thus, for the highly conserved COI protein there is a disconnect between some of the bases in its gene and the phenotype. Some bases in the gene relate to that phenotype. Others relate to the species that contains that gene.

### Coding Strategy

The genetic code is degenerate (redundant) and any one of a set of “synonymous” three-base codons can encode the same amino acid. Synonymous mutations, which change the codon without changing the amino acid, can dramatically affect GC% values; they generally locate to the third of the three positions in a codon. On the other hand, amino acid-changing (“nonsynonymous”) mutations primarily affect the second of the three codon positions. To meet its needs, each species employs some of the 64 possible codons more than others. This is its “coding strategy.” In the insightful “genome hypothesis” that provides a theoretical underpinning for much of this paper, Grantham et al. (1986)took the view “that coding strategy is a fundamental evolutionary structure and that species or kinds of species can be characterized by variation in this structure.”

A way of studying the coding strategy of different species is to plot the GC% values for each codon position against the average GC% of all three codon positions (Muto and Osawa 1987). Fig. 1 shows, for multiple bacterial species, plots of the average base composition for each codon position of the set of all the sequenced genes of a species, against the average base composition of the genes of that species (Lee et al. 2004). Low GC% species are at the left and high GC% species are at the right, and there are three data points for each species. Each codon position increases its GC% as the corresponding genomic GC% increases. The slopes of the three plots summate to 3. If a codon position were to make no contribution to the increasing GC% then its slope value would be zero. If changes in just one codon position were sufficient to change the total GC%, then its slope value would be 3. In practice we see that in bacteria, as in other groups of organisms, the major contributors to increasing species GC% values are third codon positions (slope 1.88). Amino acid-determining second codon positions that are constrained by phenotypic demands make the least contribution (slope 0.43). First positions play an intermediate role.

**Fig. 1.**
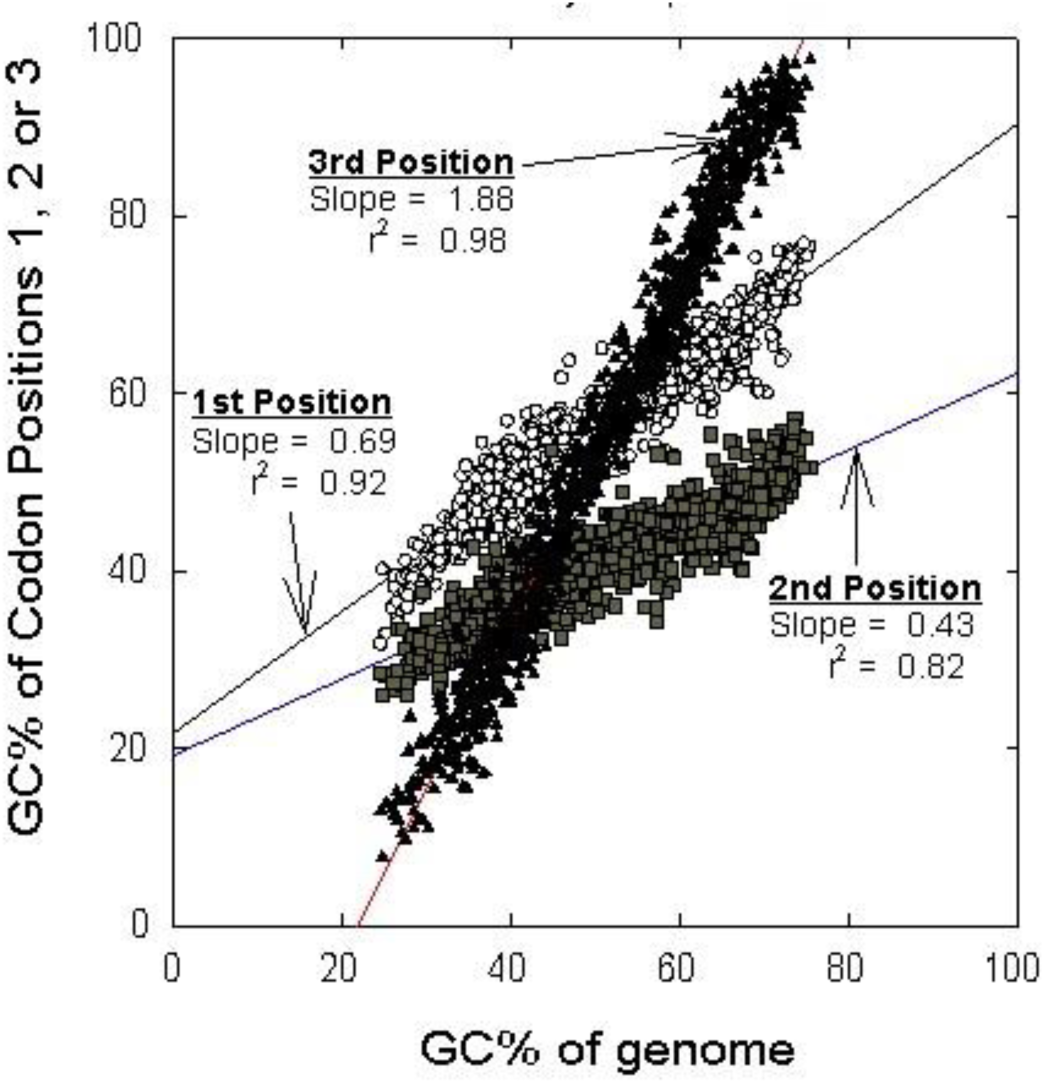
Linear regression plots of the base composition (GC%) of individual codon positions against that of all codon positions. For sequenced genes of each of 1046 bacterial species there are three values corresponding to the average GC% values of first codon positions (white circles), second codon positions (grey squares), and third codon positions (black triangles). Values for slopes and squares of correlation coefficients (r^2^) are shown for each line. Data for this figure and Table 1 are from Lee et al. (2004)and Min and Hickey (2007).

This approach allows comparison of GC% changes not only for all genes within different species as in Fig. 1, but also for a single gene within different species. For COI, whose high conservation indicates stringent phenotypic demands across a wide range of mitochondrial species, we would expect few changes in amino acids as the GC% of those species increases. Table 1 compares the Fig. 1 slope values for bacteria, with those for animal mitochondria (Min and Hickey 2007).

**Table 1.** Slopes of linear regression plots of GC% values of individual codon positions.

| **Codon position** | **1** | **2** | **3** |
| --- | --- | --- | --- |
| **1046 Bacterial species (Fig. 1)** | **0.69** | **0.43** | **1.88** |
| **849 animal mitochondrial species** | **1.01** | **0.44** | **1.55** |
| **849 mitochondrial COI subunit genes** | **0.90** | **0.09** | **2.00** |
| **4290 *E. coli* genes** | **1.00** | **0.55** | **1.45** |

For *all* mitochondrial proteins the second position slope value is, on average, 0.44, much like that of bacteria. However, in the case of the COI barcode the slope is near zero (0.09), and, while first codon positions play an intermediate role, third codon positions are essentially given free rein to generate the species-specific GC% values (slope 2.0). Thus, in keeping with its barcode role, there is large disconnect between the conventional phenotype (second position) and the contribution to mitochondrial species GC% (third position). By largely eliminating one source of GC% change across mitochondrial species, we unmask a major source, which relates to species identification. The great utility of the barcode sequence relates to the much greater disconnect of its changes from the pressures on the conventional phenotype (slope 0.09) than the changes in the general collective of mitochondrial genes (slope 0.44). The latter are similar to that for the collective of genes of an individual bacterium *(Escherichia coli*; slope 0.55).

### Codon Information on Species

Early workers in the barcode field recognized that “information is lost as one goes from DNA to protein because of the degeneracy of the genetic code.” To the extent that “information” implies biological meaning, this correctly inferred that codons potentially contain *more* biological information than is needed for amino acid-encoding (Bartlett and Davidson 1992). On the other hand, seemingly focusing on the information needed for amino acid-encoding rather than for other roles (e.g. initiating and/or sustaining species distinctiveness), Hebert et al. (2003a)considered that “nucleotide composition at third-position sites is often strongly biased (A-T in arthropods, C-G in chordates), reducing information content.”

The incorrect notion that this third position variation *reduced* biological information content was in accord with the view that GC% fluctuations reflected some inherent “directional mutation pressure” (Sueoka 1995) or “mutational bias” (Singer and Hickey 2000) of no deep biological significance – just something, essentially neutral, built into the cellular machinery. However, from an extensive study of the information content of individual codon positions in fish species, it was concluded (Ward and Holmes 2007):

> The ability of DNA barcoding to separate species relies to a very large extent on the degenerate nature of the genetic code. Amino acid sequence information has far less discriminatory power and could not differentiate the bulk of the species examined. Those amino acid residues with known critical functions are almost completely conserved over all these 388 fish species. The low rate of nonsynonymous mutations is expected for an important functional gene like cytochrome *c* oxidase, and reflects the action of strong purifying selection in conserving amino acid residues and maintaining protein function.”

Thus, by virtue of its encoding a *highly conserved* protein, the barcode segments of the mtDNAs of different species are cleansed of the differences – potential distractors – that govern their conventional phenotypes. In this respect the sequence is evolutionarily static (Forsdyke 2013). The differences that remain, largely affecting base composition at third codon positions appear to be reflective of the way organisms fundamentally differ as species (i.e. differences in their states of reproductive isolation from each other).

That species information can disconnect from phenotype is common knowledge. Domestic dogs are members of one species yet, at the behest of human breeders, they have come to display great anatomical and physiological differences that would, to the naïve observer, suggest the existence of multiple dog species. Indeed, recent studies strongly suggest that “adaptation is largely decoupled from speciation,” so “we should not expect it to be a driver of speciation” (Hedges et al. 2015; Forsdyke 2017). Furthermore, defining the “grey zone of speciation” as between 0.5-2.0% synonymous divergence, Roux et al. (2016)conclude:

> A strong and general relationship between molecular divergence and genetic isolation across a wide diversity of animals suggests that, at the genome level, speciation operates in a more or less similar fashion in distinct taxa, irrespective of biological and ecological particularities.

Much of this had been anticipated by Grantham et al. (1986):

> Another quantitative genetic difference between species is in degenerate base use. It was formerly often thought that variation in degenerate base frequencies would be a neutral phenomenon since no direct phenotypic expression results. But, this has turned out not to be so. Systematic exploitation of the codon catalogue creates genetic distances between species …. It has been shown that the greatest determinant in creating these distances is not the protein composition; instead it is the pattern of choices among degenerate bases. … Therefore, an mRNA sequence provides a better indication of the evolutionary position of a gene than does the protein sequence it codes.

The latter viewpoint was extended by Rogerson (1991)to include species-specific constraints imposed by oligonucleotide frequencies (referred to as “short sequence distribution”):

> The base sequence of the DNA … has a fundamental constraint, which underlies all other information. … These seemingly universal short sequence constraints could be contributors to many other patterns in DNA, especially the biased usage of codons within coding regions …. Codon bias in particular could be caused by the imposition of one structural design (the codons) on top of a second design (the short sequence distribution). This would rationalize the genome hypothesis of Grantham …, who showed that all of the genes in a given organism seemed to conform to a common codon usage pattern, and suggested that there might be a genomic system for selecting codons. If sequence constraints vary from species to species …, the very presence of constraints might be related to the process of speciation.

Grantham’s disparagement of neutral theory was supported by later barcode work (Stoeckle and Thaler 2014). I now turn to the questions of whether mitochondrial barcode differences could have actually driven species divergence (Lane 2009; Hill 2016) and how the barcode acquired such precision in species classification.

## 4. Speciation and Mitochondrial DNA Calibration

### Primary and Secondary Reproductive Isolation

The “spark” that initiates a new species involves a mechanism for reproductive isolation so that the process is not subverted by recombination between the nDNA genomes of diverging types - such recombination would tend to homogenize rather than retain differences. Reproductive isolation begins with interruption of the reproductive cycle - gamete, zygote, embryo, meiotic adult gonad, gamete, etc.. Being a recursive cycle, any point, be it before or after union of gametes to form a zygote, will serve to mediate the primary interruption. Whatever the point, for successful branching evolution, two independent cycles - two species - must eventually emerge.

There are many postulated mechanisms for initiating cycle interruption. If the primary interruption is at meiosis, so that gamete formation is impaired, then the parents are reproductively isolated from each other. The hybrid sterility of their *otherwise normal* offspring is a manifestation of *their* partner-specific genome phenotypes - disparate parental oligonucleotide frequencies that impair the chromosome pairing necessary for nDNA error-correction and the maintenance of species bounds (Forsdyke 1996; 2001; 2016; 2017; Reese and Forsdyke 2016). Only within the bounds of their new species will non-disparate partners be found.

This *primary* disparity – usually of non-genic origin – may serve for a while to secure cycle independence. But subsequently a *secondary* disparity may arise. Shielded by the primary barrier, members of one incipient species may accept mutations in genes such that they no longer interact properly with genes of the other incipient species. This secondary barrier between the species may affect the development of their offspring. They are *not normal* (hybrid inviability). In some circumstances such developmental failure may also be a primary cause of cycle interruption (Fig. 2).

**Fig. 2.**
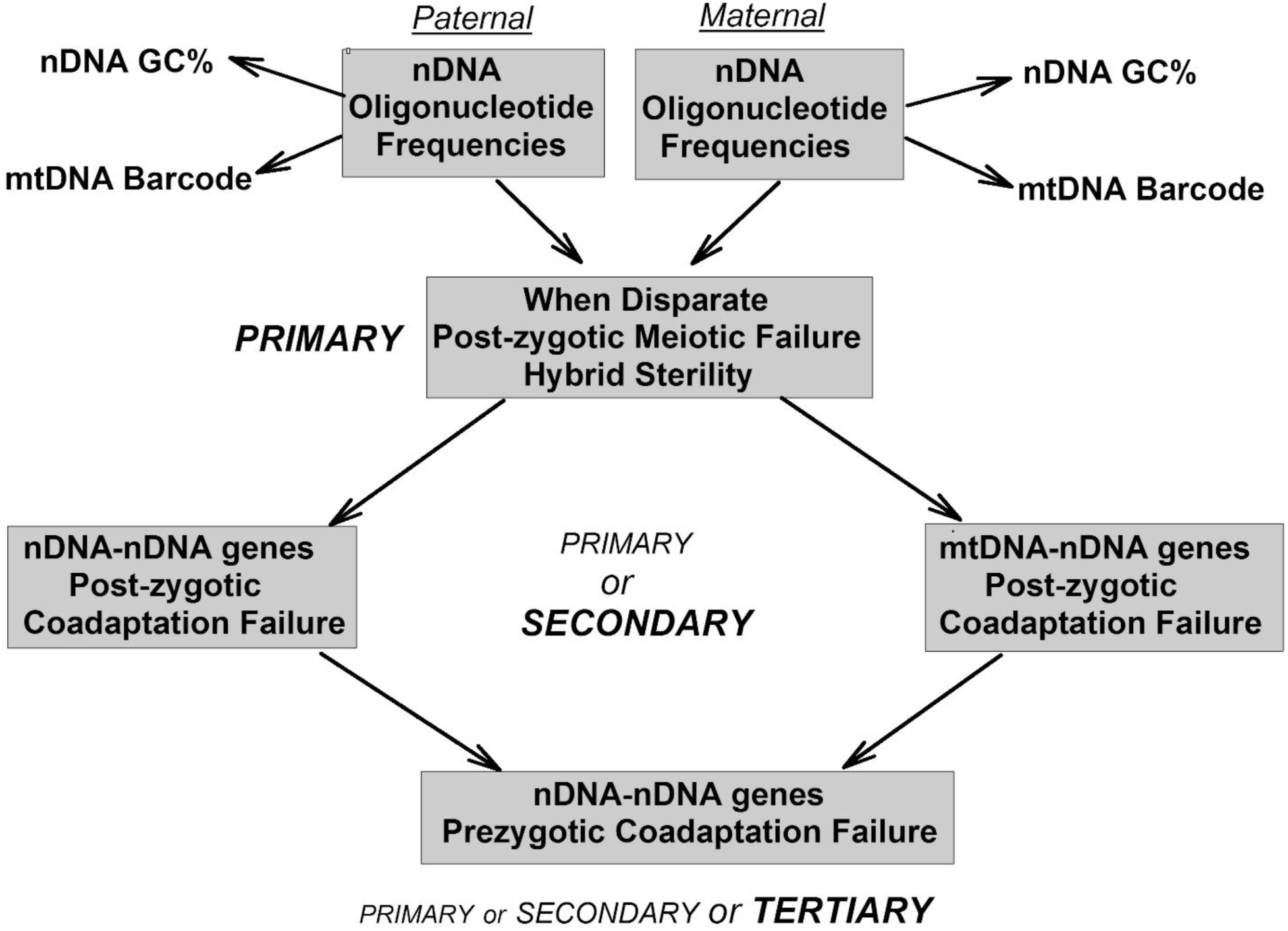
Alternative paths for primary initiation of reproductive isolation that can lead to divergence into two species. Most modern species are at the prezygotic tertiary stage (bottom) where multiple genes fail to coadapt (e.g. mouse cannot copulate with elephant). Although initiation can occur at any level, the general primary cause (indicated by size of italic capitals) is here held to be the hybrid sterility of otherwise normal offspring (top). This is a manifestation of disparate parental oligonucleotide frequencies that impair meiotic recombination. While failed coadaptation of mtDNA-nDNA gene products (right) is here secondary to a prior initiation (top), some emphasize the possibility that it may be primary (see text). Unrelated to speciation mechanisms, the mtDNA barcode phenomenon (top left and right) is a species specific expression of parental nDNA oligonucleotide frequencies, much as is the, albeit less specific, nDNA GC%.

Thus hybrid inviability can be either primary or secondary. However, hybrid sterility can *only* be primary, since if a hybrid is inviable then there can be no adult hybrid to manifest sterility.*The onus is upon those believing a case of hybrid inviability to be primary, to show that it had not been preceded by a period, however brief, of hybrid sterility.*

### Potential Genic Conflict

There may be primary or secondary *genic* sources of cycle interruption. Gershoni et al. (2009)suggested that among the “alternative mechanisms to drive speciation events” could be incompatibility between parental mitochondrial and nuclear genomes. In their view, Herbert’s *“pas de deux* of surprising intimacy” that links mitochondrial and nuclear genomes largely reflects the *coevolution* of certain genes and hence of the corresponding gene products. If coevolution fails there is a potential for genic conflict. Each new generation the mitochondrial genomes contributed unilaterally by the maternal ovum are forced to “reassess” their relationship with both the novel paternal nuclear genome that is encountered and the maternal meiotic products co-contributed by that ovum.

This is a two way process. Mitochondrial and nuclear genomes have to *coadapt.* In vertebrates, the nDNA has many genes (~1500) that encode proteins destined to function in mitochondria. The mtDNA itself has only 13 protein-encoding genes whose products normally function in the mitochondrion where they are synthesized. One is the “sentinel” barcode gene, cytochrome c oxidase subunit I (COI), which contains the 648 base barcode sequence. COI is a subunit of the multisubunit cytochrome oxidase enzyme, 10 subunits of which are encoded in the nucleus. A requirement for *physical interaction* between subunits encoded by nDNA and mtDNA has long been known. Schmidt et al. (2001)concluded for cytochrome oxidase that: “the faster mtDNA mutation rate … makes mtDNA the predominant partner in accommodating mutations important for subunit interaction.”

Thus, benefitting from this high mutation rate, an amino acid change in a paternal nDNA-encoded mitochondrial protein would favor (create a selection pressure for) cells where some mitochondria had rapidly acquired compensatory mutations in maternal mtDNA (Hill 2016). Cells with mitochondria that did not acquire the right mutations could die, leading to the possible death of the host organism. Conversely, cells with mitochondria that had acquired the right mutations would thrive, but could only be transferred successfully to future host organisms with compatible nDNA. Indeed, keeping up with the pace set by the mitochondrial genome, some nDNA-encoded subunits involved in the mitochondrial electron-transport chain can display increased mutation rates that are consistent with continuous positive selection for compatibility (Mishmar et al. 2006; Gershoni et al. 2010; Bar-Yaacov et al. 2015).

So mtDNA can respond to (track) nDNA and, at the outset, Hebert et al. (2003b)envisaged “the rise of mitochondrial variants with more effective nuclear interactions.” However, as they say, the devil is in the details, and the onus is on those advocating mitonuclear *genic* conflicts as explanations for the barcode’s taxonomic power and its possible general role in speciation (Lane 2009; Hill 2016), to provide those details. Although much work remains, the following *nongenic* hypothesis might meet this need.

### How mtDNA Might Monitor nDNA Oligonucleotides

How do genome-wide oligonucleotide changes in nDNA, which are reflected in a distinctive nDNA GC% and can spark a primary speciation event, become registered as changes in mtDNA that deeply involve the COI-encoding sequence, so perfecting its barcode role? While there is much mitonuclear cross-talk (Ryan and Hoogenraad 2007; Hill 2016), less attention has been given to the information flow from nucleus to mitochondrion by way of the nDNA-encoded proteins that are synthesized in the cytoplasm, but are destined to operate in the mitochondrion.

The distinctive nDNA GC% will have *marked* the amino acid composition of those proteins (see above). How could a mitochondrion “read” the changed amino acid composition to “deduce” nDNA GC% (oligonucleotide frequencies) and hence the species within which it resides? Why is it that the COI-encoding sequence is so well fitted to “speak for” the mitochondrion in this respect?

A clue here is that, in contrast to most organisms, the mitochondrial repertoires of the RNAs that transfer amino acids to the ribosomes for protein synthesis are limited. The number of transfer RNAs (tRNAs) matches the number of amino acids (20), plus two extra tRNAs for the amino acids (arginine and serine) that normally have six codons instead of the usual four. Thus, for mitochondrial protein synthesis there is basically only *one* tRNA for each amino acid. While there is some flexibility in that one tRNA can sometimes recognize (though not with equal accuracy or efficiency) more than one codon corresponding to a particular amino acid, for the usual four-codon amino acid three synonymous codons in mitochondrial mRNAs may be underemployed.

The nature of the underemployed codons varies with the species. Mitochondria from different species have their own bias as to which synonymous codons to “deem” optimum and which to under-employ. Assuming the option of increasing tRNA number is not available, a way this situation can change as species change is by mutational conversion of some tRNAs to those better recognizing one of their previously underemployed synonymous codons. This mutation will usually be in the base that matches the third codon position in the mRNA that is being decoded. If a blanket change in certain tRNAs suffices to bring all mRNAs into line there is no need for the mRNAs themselves to adapt at multiple codon positions. If the mitochondrion in which a tRNA mutation occurs thrives, then its frequency within the mitochondrial population within its host cells should increase, as will the mutation it contains.

A second clue is the large number of nDNA-encoded proteins (~1500) that are destined to function in mitochondria. The changes in nDNA GC% levels that relate to the species-initiation mechanism discussed above, will, by affecting first and second codon positions, have changed some of the encoded amino acids in proteins that are not under strong selective constraint. In turn, some of the small subset of thirteen mitochondria-encoded proteins, which interact with some of these nDNA-encoded proteins, may need to change their properties either to sustain this interaction, or prevent adverse interaction. This may make it advantageous for the corresponding mitochondrial genes, *if not highly conserved* through other selection pressures, to accept mutations that affect first and second codon position, so changing the nature of the mtDNA encoded amino acids.

Changes in the usage of the *single* tRNAs corresponding to each of these changed amino acids (increases or decreases) might be accommodated. But it is likely that for some mitochondrial amino acids a particular synonymous tRNA will work more accurately or efficiently (Dana and Tuller 2014; Du et al. 2017). Thus, there will be an evolutionary selection pressure on some of the 22 mitochondrial tRNA genes to accept mutations that change the synonymous codons to ones that best meet the new challenge. This, in turn, will pressure a *highly conserved* gene, such as COI, to adapt its third (and to some extent first) codon positions.

Thus, by this tortuous path (Fig. 3) COI, while rigidly retaining its second codon position fidelity, will through changes in other codon positions to better match the changed mitochondrial tRNA population, accurately monitor the GC% changes in the corresponding nDNA. That this scheme could explain Herbert’s “ferocity of lineage pruning” is suggested by the growing recognition in other systems that “evolutionary shifts in the identity of selectively favored codons can occur” so that “the identity of favored codons tracks the GC content of the genomes” (Hershberg and Petrov 2009; Forsdyke 2016, p. 209-234). Correlations between mtDNA barcodes and the corresponding mitochondrial tRNA repertoires might provide useful support. Evidence that “mtDNA substitution rates in a lineage are positively correlated with nuclear DNA substitution rates in the same lineage” (Eo and DeWoody 2010) is also supportive.

**Fig. 3.**
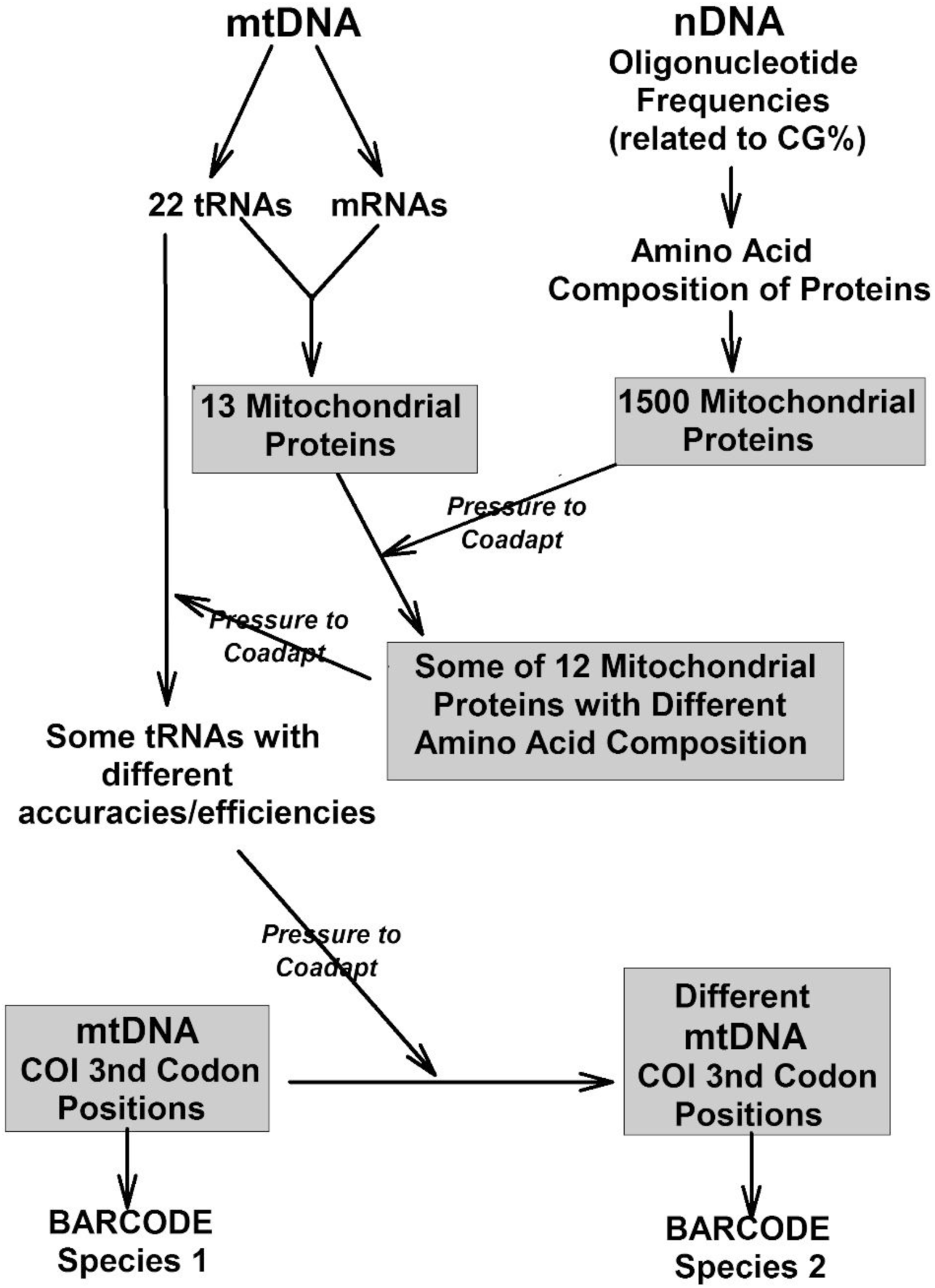
How changes in nDNA oligonucleotide frequencies that initiate species divergence may, in a short evolutionary period, be registered in the sequence of the most conserved of the thirteen mitochondrial protein-encoding genes (barcode sequence). Coadaptive pressures pass from (i) nDNA-encoded mitochondrial proteins to some of the less-conserved mtDNA-encoded mitochondrial proteins with which they interact, then on to (ii) mitochondrial tRNAs that decode the mRNAs for those mitochondrial proteins, and finally to (iii) those codon bases in the most-conserved protein-encoding mtDNA gene that do not determine amino acid content. For details see text.

## Conclusion

Hebert envisaged a “scouring mechanism” by which the within-species diversities of mtDNAs are decreased, so facilitating fine barcode discrimination between species. This depends on the ability of the rapidly-mutating, non-amino acid-determining, nucleotide bases in mtDNA codons to track the more stable, but varying, oligonucleotide frequencies of the different nDNAs that mitochondria must coexist with, generation after generation, within a common cell. Mutational fluctuations of nDNA oligonucleotide frequencies around the norm for a species – long ago detected by researchers as genome-wide changes in nDNA GC% - impair the meiotic pairing of chromosomes, so initiating the reproductive isolation necessary for sympatric divergence into two species. Some nDNA oligonucleotide changes involve second codon positions so affecting the amino acid composition of proteins and changing their properties, which include interacting with other proteins.

Changed proteins include the ~1500 that are normally transferred to the mitochondria where some interact with some of the 13 proteins that are locally encoded by mitochondrial genes. Those mitochondrial genes that encode *less conserved* proteins have sufficient flexibility to accept mutations in second codon positions that change their amino acids, so improving the interaction. As a byproduct of this, the mitochondrial tRNA repertoire is modified through mutations in tRNA genes to facilitate more accurate and efficient recognition of codons. The *highly conserved* COI gene tracks these newly favored codons mainly by accepting *synonymous* mutations. Since non-synonymous amino acid-changing mutations are infrequently accepted, the species individuality of each COI gene is an unhindered expression of the individuality of the species itself as first determined by its nDNA oligonucleotide frequencies.

A need to constantly calibrate mtDNA against nDNA would seem to explain why the disparate organelle and nuclear genomes are linked in an intimate *pas de deux,* and to resolve the mystery of the mechanism and biological significance of the barcode phenomenon. A better understanding of why the COI barcode mechanism works so well in animals could assist the search for barcodes in plants where the COI barcode is less successful. There are also implications for the emerging field of mitochondrial replacement therapy in humans (Latorre-Pellicer et al. 2016).

## Acknowledgements

Queen’s University hosts my web pages that display some of the early speciation literature (http://post.queensu.ca/~forsdyke/evolutio.htm or https://archive-it.org/collections/7641).

